# Proliferation in the epiblast-hypoblast layers of the primitive-streak chick embryoblast in response to epidermal growth factor

**DOI:** 10.1101/706630

**Authors:** Francis O. Cunningham

## Abstract

The reaction of twenty four hour old primitive streak chick embryoblasts exposed to epidermal growth factor was compared with non-exposed controls. After 24-36 hours incubation, proliferation up to 5-6 cell layers in thickness of the epiblast-hypoblast layers was evident in the EGF exposed embryo blasts. This compared with the normally expected one cell layer in thickness at this stage of development.

## Introduction

Epidermal growth factor (EGF) was discovered over 60 years ago during a search for nerve growth factor. Cohen demonstrated that extract from mouse submaxillary gland stimulated epithelial cells derived from the dorsal skin of the 7 day old chick embryo (Cohen,1965). There was marked increase in the number of epidermal cell layers in the dorsal skin exposed to the mouse extract. Similar mouse extract induced precocious eyelid opening and incisor teeth eruption when injected into new-born animals by stimulating epidermal growth and keratinisation. The factor causing these effects was isolated and found to be a low molecular weight, heat stable non-dialyzable polypeptide. It was called epidermal growth factor (Cohen,1983). Cohen concluded that EGF stimulated growth factor receptors (EGFRs) in the seven day old chick embryo dorsal skin. The question as to how early in embryonic development that EGF receptors are present was unknown until reports of implantation of non-human cancer cells in the chick embryoblast (Lakshmi and Sherbet, 1974),(Palayoor and Batra, 1971). Later, implants of human malignant cells, placed between the epiblast-hypoblast layers in primitive-streak chick embryos, were shown to induce ectodermal (ECTDPR) and endodermal proliferative(ENDPR) responses (Cunningham, 2019). These observations suggested that malignant tumour cells were secreting growth factors (EGF,TGF etc.) which reacted with receptors in the primitive streak stage epiblast-hypoblast layers.

### Aims of the study

We studied the reaction of the 24 hour old primitive streak epiblast-hypoblast layers in the chick embryoblast exposed to EGF.

## Materials and Methods

A standard bacteriological incubator was used for incubating the eggs and explanted embryoblasts and tap water provided humidity at 60%.

Heat resistant Pyrex glassware and an aluminium egg separator were sterilised at 200 deg. overnight.

An ultra violet light cabinet provided a sterile environment for the cooling glassware, in particular a large open Pyrex^R^ dish.

A Nikon SMZ10 ^R^ dissecting microscope facilitated implantation of cells in the embryoblast. Steriised J cloth was cut into rings to fit inside the Petri dish underneath the watch glass. (Johnson & Johnson^R^).

Freshly fertilised broiler hens’ eggs (Gallus Domesticus) of Arbour Acres breed were obtained twice weekly and stored at room temperature until placed in the incubator. Eggs were used within three days of receipt and the fertility rate varied between 80 to 90 %. The use of chick embryoblasts for research was in accordance with national and international guidelines.

Pyrex glass tubing with 1mm wall thickness and 10mm inside diameter, was cut in 5mm. lengths to give glass rings to hold the vitelline membrane in place on a watch glass. A fine stainless steel probe was made by cutting a 3 cm. length of 0.035mm wire and placed in a glass Pasteur pipette and flamed into position. The protruding wire, cut obliquely to a length of 0.5cm., was used to incise the epiblast.

Defined Medium 199 was used for transport of the cells or tissue under study. (Flow Laboratories England, supplied by Medlabs Ltd. Ireland).

Dulbecco ‘A’ solution was used to float the egg yolk and harvest the vitelline membrane with the attached primitive streak. (Oxoid Ltd).

Pannett-Compton solution, the supernatant left over after mixing Dulbecco ‘A’ and ‘B’ (Ca++,Mg++ salt solution), was used to cover the embryoblast in culture.

Natural medium thin egg white albumen, the nutrient medium of the embryoblasts, was recovered after separation of albumen and the egg yolk in the sterilised aluminium separator. An essential growth stimulator, its colloid osmotic pressure is of great physiological importance for the exchange of water in the embryo (Schmidt, 1993).

Epidermal growth factor 100ug. was obtained from the Sigma Chemical Company (supplied by Med Labs. Ireland. EGF No. E-7755 from mouse submaxillary glands, sterile filtered). This was dissolved in 10 ml. of Pannett Compton solution under sterile conditions. Aliquots of 1 ml. (10 ug) were stored at −20 °C until further use.

### Explantation Technique

Egg viability was confirmed by ‘candling’. This was done by trans-illuminating the egg sitting on a holder with a bright light underneath. The viable embryo was seen as a dark spot inside the shell.

Two dozen eggs were incubated at 37 ° C. for 20-22 hours at 60 % humidity and then left at room temperature for 1-2 hours before use. Each egg in turn was cracked open onto the sterile aluminium egg separator to remove the albumen. The thick albumen in the beaker below was poured off to leave the thin albumen. One could tell on gross examination that there was a viable embryoblast. The egg yolk was floated in a bath of saline and the vitelline membrane was cut round its equator with an iris scissors. The vitelline membrane was peeled off the egg yolk with two forceps and floated onto a watch glass and covered with a pyrex glass ring. The watch glass and ring were removed and placed in a Petri dish on top of the central hole in the sterile circle of J cloth. The dish was placed on the stage of the dissecting microscope. Light from below was transmitted to the embryo, through the hole in the J cloth. The vitelline membrane edge was draped over the glass ring from outside to in. The saline inside and outside the glass ring was pipetted away. This prevented the whole assembly from drifting on the watch glass. The free edge of the vitelline membrane was now pulled taut and even over the upper edge of the glass ring until the membrane was flat with the embryoblast at the centre of the ring. Any wrinkles at this stage were ignored as they disappeared as the embryo was cultured. The edge of the vitelline membrane inside the ring was then trimmed. This facilitated removal at the end of culture.

With the aid of the dissecting microscope, the primitive streak was easily recognised. The fine wire probe was flamed and an incision was made in the epiblast at the edge of the area pellucida above the level of Henson’s node separating epiblast-hypoblast layers. Twenty explanted embryoblasts were used. The epiblast-hypoblast layers were separated as far as the primitive streak in one half of each embryoblast. This exposed the epiblast to the nutrient PC solution. Ten of the embyroblasts had Panett-Compton (PC) solution 0.3ml. only pipetted inside the ring (controls), while ten more had PC with EGF (3ug.). Thin egg albumen 0.6 ml. was pipetted outside the ring on the watch glass. The J cloth was saturated with saline to provide a moist chamber. They were incubated for 20-22 hours.

### Fixation and processing of specimens

After fixation in Bouin’s fluid, specimens were floated in formalin on a flat piece of paper and placed in a specimen holder for identification in an automatic tissue processor (Shandon^R^). The specimens were cut square across at the cephalic end and cut to a point at the tail end for orientation before embedding in wax. This aided recognising the cephalic end as the specimen was very small and showed as a small streak of yellow in the block. The wax around it was trimmed carefully so that only a small amount surrounded the specimen. It was sectioned from head to tail by the microtome. Serial ribboned 8u thick sections, placed in sequence in two rows of 25-30 sections on each slide, fitted a 400-500u piece of embryo on each glass slide. Each sectioned embryonic specimen fitted on 5 to 6 slides. Stained with haemotoxylin and eosin, sections were studied in sequence from head to tail under the microscope looking for proliferative reactions.

## Results

Twenty four embryos were explanted and 16 were grown to 14 somite stage with normal development. Three of eight controls grown without EGF were lost in processing. Eight of 8 embryos treated with EGF showed ECTPR and ENDPR. Some showed an organised epithelial appendage.

**Figure A.**
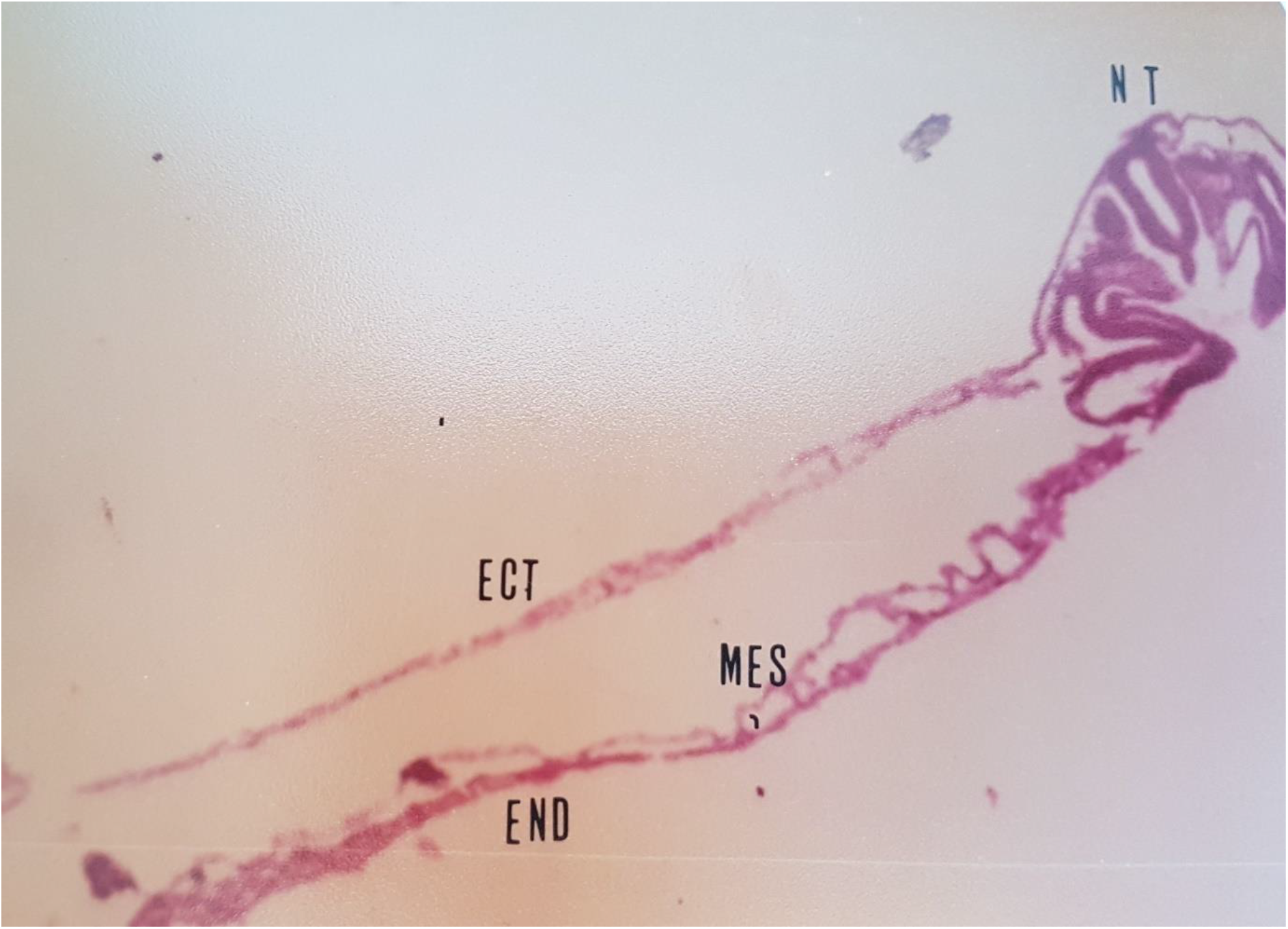
One to two cell layers of normal ectodermal (ECT), mesodermal (MES) and endodermal (END) development with neural tube (NT).

**Figure B.**
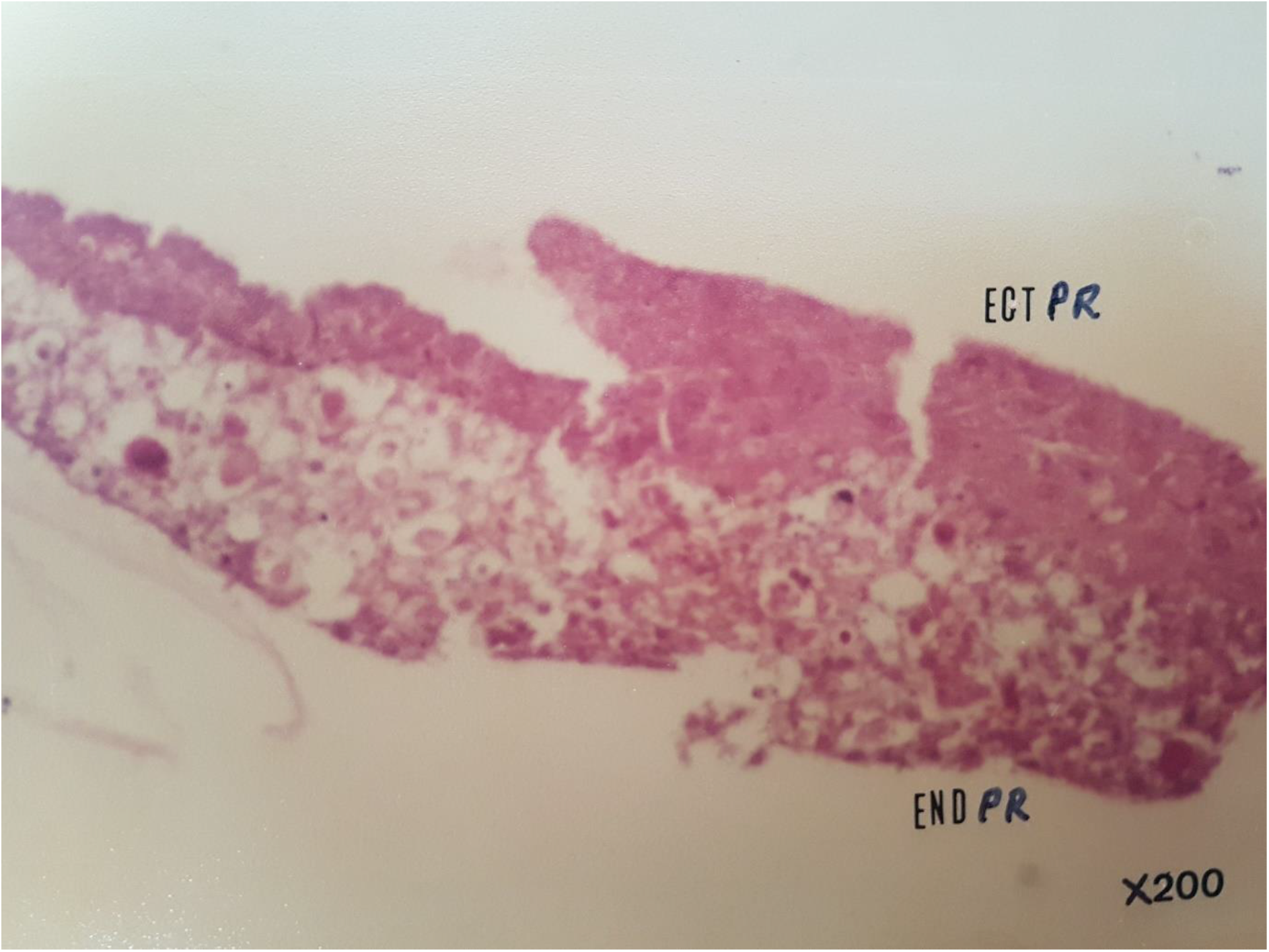
Ectodermal ENDPR) and Endodermal (ENDPR) proliferation in response to EGF.

**Figure Ca.**
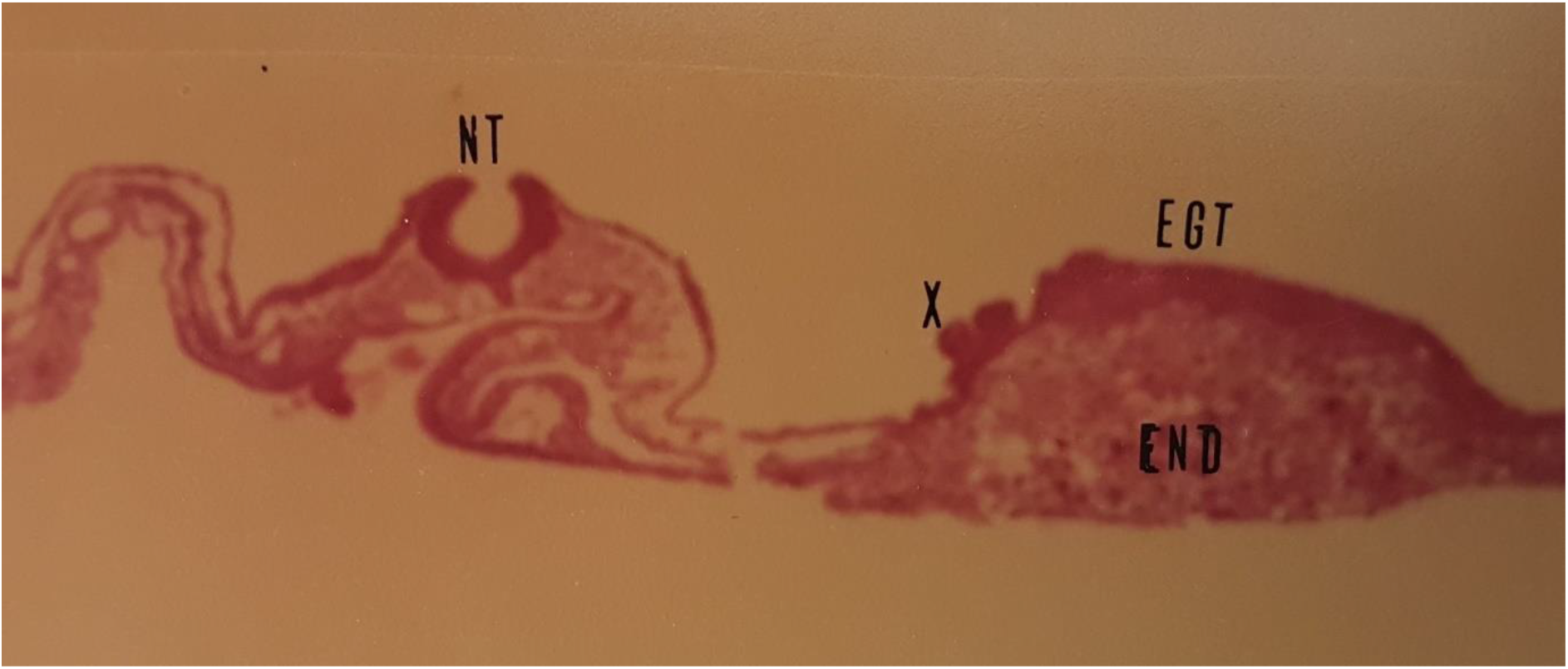
shows ECTPR,ENDPR reactions to EGF. Neural tube (NT).

**Figure Cb.**
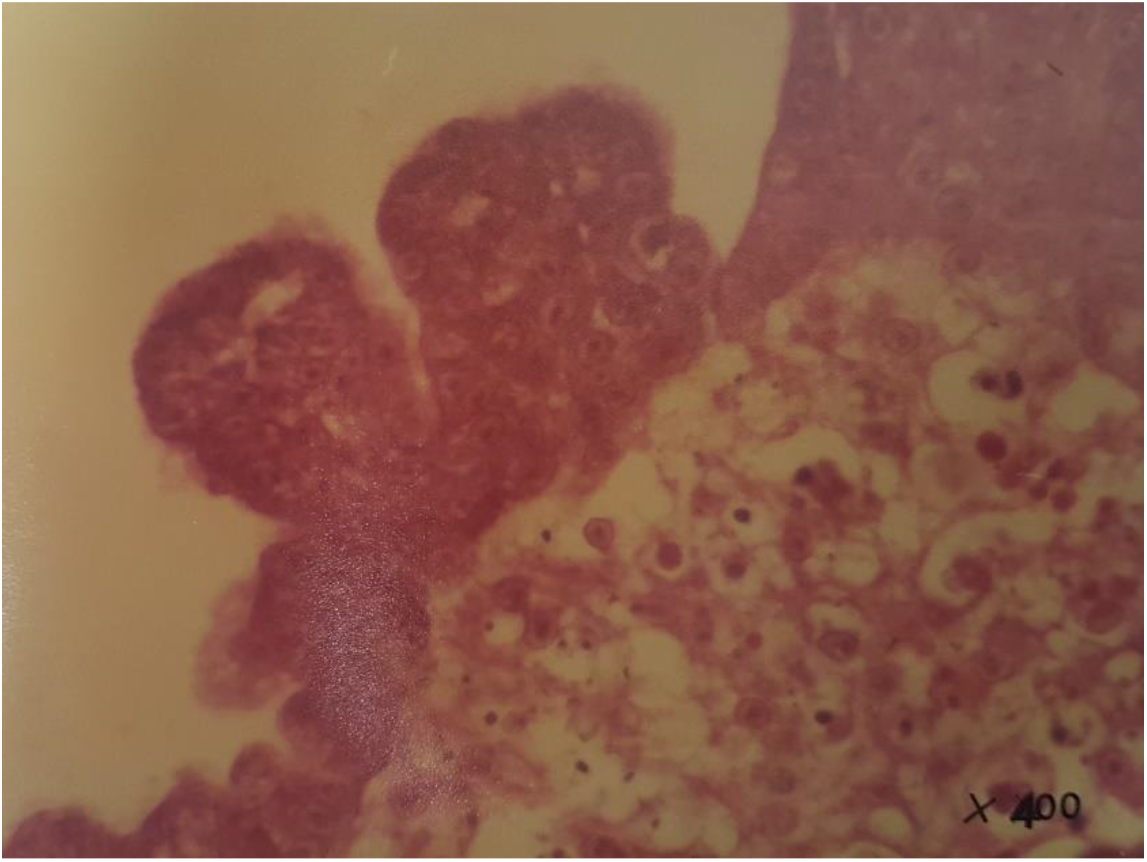
Area X of Ca magnified in Cb.

**Figure D.**
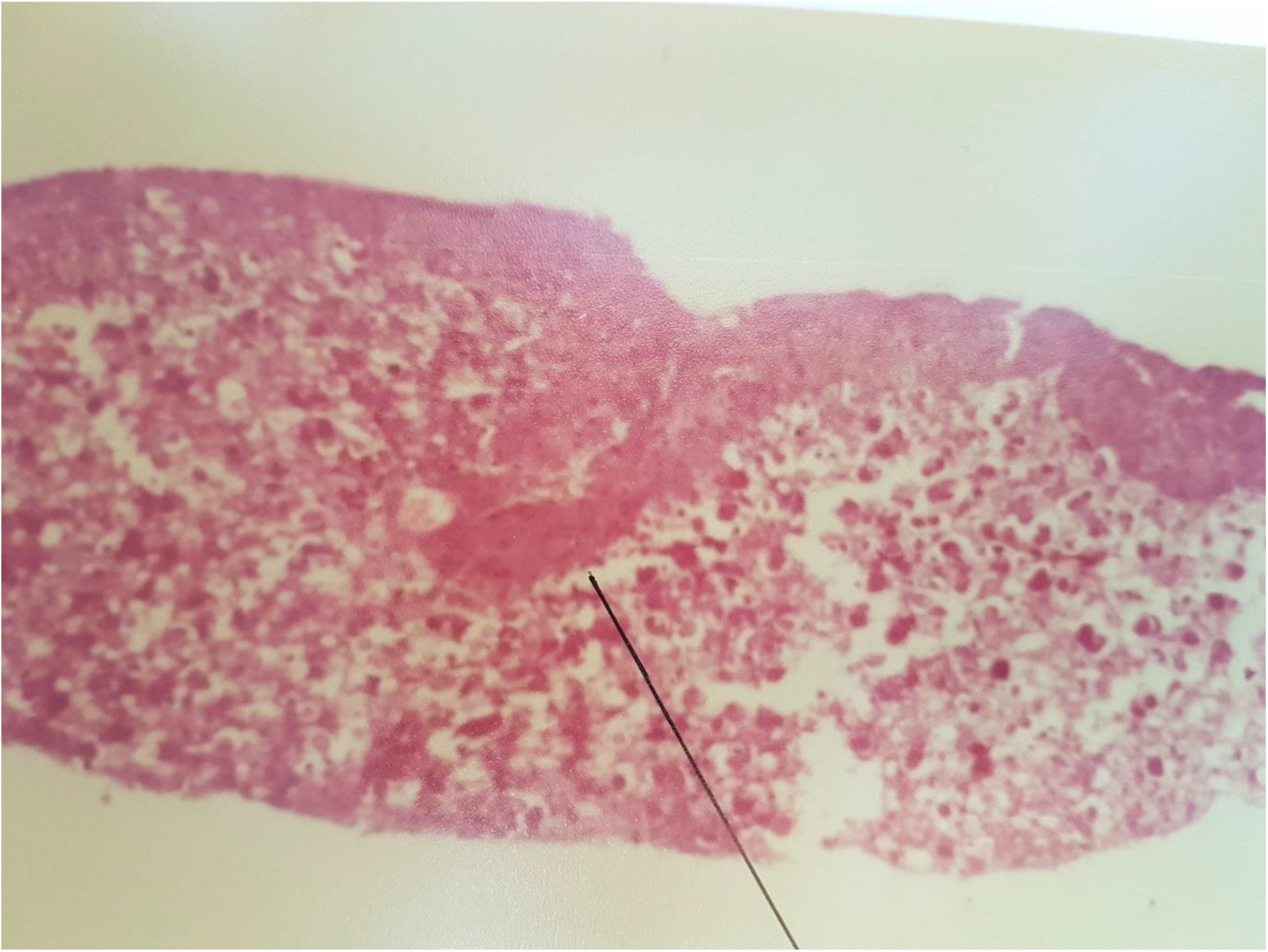
Shows ECTPR with organised ectodermal structure in EGF treated embryoblast, black line.

## Discussion

Cohen’s discovery of the proliferative reactivity of the dorsal skin of the 7 day old chick embryo to mouse sub-maxillary gland extract, eventually led to the discovery of EGF. This implied that the primitive skin had EGF receptors. It was not known how early in development these receptors appeared in the embryoblast until non-human cancer cells were implanted by Sherbet and Lakshmi into primitive streak chick embryoblasts giving a proliferative reaction (Lakshmi and Sherbet,1974). This was confirmed by Palayoor and Batra (1971). More recently, the first report of implants of human cancer cells from malignant breast, colonic and rectal tissues, human colonic cell line (HT-29) and a nasal squamous cell carcinoma cell line (RPMI-2650) into primitive streak stage chick embryoblasts, showed that they provoked proliferative reactions ECTPR and ENDPR (Cunningham, 2019).

This study confirms that the proliferative reactions of the chick primitive streak layers to implanted human cancer cells and exposure to EGF are similar. The presence of EGF receptors in primitive streak stage chick embryoblast layers is demonstrated for the first time. Histological EGFR staining of the primitive-streak stage embryoblasts is the next step to see the extent of expression of these receptors.

## Abbreviations

ECTPR: ectodermal proliferative response
HMPR: mesodermal proliferative response
ENDPR: endodermal proliferative response
EGF: epidermal growth factor EGF
TGF: transforming growth factor

## Acknowledgements

I am grateful to Professor Niall O’Higgins St. Vncent’s Hospital, Dublin, who suggested this research. Dr. G. Sherbet and Dr. Lakshmi kindly taught me the explantation technique at Newcastle-Upon Tyne. Their helpful letters steered me through my initial errors. Invaluable at Woodview were Nicholas O’ Connor, Chief supervisor; Dr.Mary Sharp for all things biochemical; Geraldine O’Neill for day to day running of experiments; Erwin Hubrich for manufacture of the the U.V. cabinet, pyrex rings and incubator box. I wish to thank Rosaleen Rafferty at St. Vincent’s Pathology Department for mounting and processing specimens, Derek Cullen at St.James’ Hospital laboratory for processing the histology. At University College Galway, Professor Sean O’Beirn and Professor H.F. Given were supporters with Michael Maguire and Tom Rogers in the pathology department. The Cancer Research Board of the Irish Cancer Society gave me a grant for equipment and running costs for the first year.

